# Senescent fibroblasts interfere with the tumor immune response in humanized models by inducing neutrophil extracellular traps

**DOI:** 10.64898/2026.02.09.704891

**Authors:** Monica Cruz-Barrera, Anthony Sonn, Cristian Camilo Galindo, Georgio Mansour Nehmo, Oanh Le, Gerardo Ferbeyre, Christian Beauséjour

**Affiliations:** Centre de recherche du CHU Sainte-Justine, Montréal, Québec, Canada; Département de pharmacologie et physiologie, Université de Montréal, Montréal, Québec, Canada; Division of Clinical and Translational Research, McGill University, Montréal, Québec, Canada; Département de Biochimie et Médecine Moléculaire, Université de Montréal, Montréal, QC H3C 3J7, Canada; Centre de Recherche du Centre, Hospitalier de l’Université de Montréal (CRCHUM), Montréal, QC H2X 0A9, Canada

## Abstract

Neutrophils are known to promote tumor progression and metastasis, in part through the release of extracellular traps (NETs), a phenomenon associated with poor prognosis in patients. Here, we demonstrate that senescent fibroblasts, via their senescence-associated secretory phenotype (SASP), promote the recruitment of CD33^+^ myeloid cells and impair the tumor immune response in an orthotopic humanized mouse lung cancer model. Mechanistically, we found that therapy-induced senescence triggers the formation of NETs, which interfere with immune cell infiltration in tumor spheroids. This phenotype was reversible upon treatment with DNAse I or Reparixin, an inhibitor of the action of CXCL8/IL-8 and CXCL1/GRO1, two key SASP factors. Furthermore, we show that senescence-induced NETs favor lung metastasis in a humanized mouse model, a phenotype that was inhibited by DNase I treatment. Our findings provide critical insights into the complex interplay between cellular senescence, NETosis, and the tumor immune response, highlighting another mechanism by which senotherapies can improve cancer treatments.

## INTRODUCTION

Neutrophils are the most abundant white blood cell type, comprising 30% to 70% of total leukocytes (1, 2). Upon activation, neutrophils can eliminate pathogens or abnormal cells through mechanisms such as phagocytosis, degranulation, and the release of neutrophil extracellular traps (NETs) (3). While neutrophils play a pivotal role in innate host defense against infections, they have also been implicated in a pro-tumorigenic role. Neutrophils promote tumor progression by inducing epithelial-to-mesenchymal transition, stimulating angiogenesis, remodeling the extracellular matrix, and suppressing T cell-dependent antitumor immunity (2, 4–7). Consequently, increased neutrophil recruitment has been associated with poorer survival outcomes in patients with solid tumors (8, 9).

In the tumor microenvironment (TME), chemokines activate neutrophils, inducing a form of cell death known as NETosis. This process is characterized by the release of NETs, which consist of decondensed DNA in complex with histones and granule proteins such as neutrophil elastase (NE) and myeloperoxidase (MPO) (10–12). Beyond their role in tumor progression and metastasis, NETs have been shown to inhibit the efficacy of cancer immunotherapy. This occurs via the physical coating of tumor cells, which hinders both their recognition by cytotoxic immune cells, including CD8⁺ T cells and NK cells, and the infiltration of these effector populations (11). These findings highlight the critical role of NETs in shaping the TME and modulating immune responses (13).

In parallel, cellular senescence is another important phenotype that has been shown to promote tumor growth and impair immunotherapy in mice (14, 15). Therapy-induced senescence is often an inevitable consequence of cancer treatments in patients (16). Senescent cells secrete a variety of cytokines, chemokines, growth factors, and extracellular matrix components, collectively known as the senescence-associated secretory phenotype (SASP) (17, 18). Several studies have demonstrated that chemokines such as CXCL8/IL-8, CXCL1/Gro-α, and CXCL2/Gro-β act through the CXCR1 and CXCR2 chemokine receptors on neutrophils, thereby modulating NET formation (11). Interestingly, these chemokines have also been identified as components of the SASP (18, 19).

To explore the interplay between the SASP and NETosis in the context of cancer therapy, we utilized humanized mouse models where tumors can be rejected following the injection of autologous human immune cells (20). Our results showed that senescent human dermal fibroblasts (HDF) promote myeloid cell recruitment, lung epithelial tumor cell growth, and metastasis. We further demonstrated that HDF, through their SASP, induce the formation of NETs, which lead to impaired tumor immune cell infiltration. This phenotype was prevented by inhibiting NET formation. Furthermore, our analysis of published single-neutrophil transcriptomic datasets (21), from colon and lung cancer patients revealed post-treatment enrichment of NETosis pathways in neutrophil subpopulations. In summary, this study allowed us to unveil the intricate interactions between cellular senescence and the formation of NETs in their roles in the cancer immune response.

## METHODS

### Cell culture

Human dermal fibroblasts (HDF) and peripheral blood mononuclear cells (PBMCs) were collected from adult donors as previously described and in accordance with the ethics committee from the Centre Hospitalier Universitaire Sainte-Justine (protocol 2017-1476) (20, 22). In brief, a skin biopsy from a healthy adult male donor (41 years old) was collected and digested with collagenase D (Roche) for 1 h at 37°C with agitation, centrifuged at 400 × *g* for 5 min, and washed with DMEM (Wisent Bio Products). HDF were cultured in DMEM with 10% FBS and 0.2% primocin. Cellular senescence was induced after treatment with 0.1µM of Doxorubicin, exposure to ionizing radiation (IR) at a dose of 12Gy (1 Gy/min using a Faxitron CP-160), or after transduction with lentiviral particles carrying an inducible version of the KRAS^v12^ gene, hereafter as RAS.

iPSCs reprogramming and cellular transformation were performed as described earlier (20). In brief, iPSCs were first differentiated into lung epithelial progenitor cells (LEC), which were then sequentially transduced with 4 distinct lentiviral particles. First, LEC were transduced with viral particles containing the SV40 large T antigen and the neomycin resistance gene. Cells were selected with 300 μg/ml G418 (Thermo Fisher Scientific) for three days, and surviving cells were then transduced with lentiviral particles carrying the HRAS^V12^ and puromycin-resistance genes. Cells were selected with 2 μg/mL of puromycin (Thermo Fisher Scientific) and then transduced with lentiviral particles carrying the catalytic subunit of the human telomerase gene (hTERT). Finally, transformed cells (LEC-4T) were transduced to express mPlum and luciferase. All transductions were carried out with 8 μg/ml Polybrene (Sigma-Aldrich).

### Conditioned cell culture media preparation and cytokine quantification

Conditioned media (CM) was collected, filtered, and centrifuged from non-senescent or senescent HDF cultured in a monolayer with RPMI 1640 (Wisent Bio Products) without serum for 24 hours. HDF were counted at the time of collection, and the CM was normalized to the lowest cell count using fresh RPMI. CM was filtered through a 0.22 μM filter to remove cell debris before being flash-frozen on dry ice. Samples were collected 8 days after senescence induction and sent for multiplex cytokine analysis (Eve Technologies, Calgary, AB, Canada) using the Human Cytokine/Chemokine Panel A 48-Plex Discovery Assay.

### Animals and tumor models

Experiments with mice were conducted in accordance with our protocol, approved by the institutional animal research committee (2022–3508). NSG-SGM3 mice, which express human IL3, GM-CSF, and SCF were obtained from The Jackson Laboratory and bred on site. Eight- to ten-week-old female and male mice were housed in controlled, pathogen-free conditions and handled with aseptic techniques. Mice were anesthetized with 2% isoflurane for cell injections and imaging.

Non-senescent or senescent HDF cells were stained with the fluorescent dye CellBrite NIR790 Cytoplasmic Membrane Dye (catalogue 30079; Biotium) following the manufacturer’s instructions before being injected into mice. 4x10^5^ HDF were injected intravenously (i.v) followed by 1x10^5^ tumor cells (LEC-4T or A549) the day after. For injection, cells were resuspended in 100 μl of RPMI 1640 (Wisent Bio Products). For subcutaneous injections (s.c.), mice flanks were shaved and 5x10^4^ tumor cells (LEC-4T) alone or mixed with 2x10^5^ non-senescent or senescent HDF were injected in 100 μl of RPMI 1640 (Wisent Bio Products). Where indicated, mice received daily intramuscular injections (i.m.) of 2.5 mg/kg DNase I (Roche Diagnostics) diluted in phosphate-buffered saline (PBS).

The growth of luciferase or mPLum-expressing tumors was monitored using the Q-Lumi In Vivo imaging system (MediLumine, Montreal, QC, Canada). Luciferase-expressing tumors were imaged 10 minutes after the i.p. injection of D-luciferin (Perkin Elmer XenoLight catalog #122799) (150 mg/kg). mPlum expressing subcutaneous tumors were tracked using the 562-40 nm excitation and 641-75 nm emission filters. Finally, NIR790-stained HDF were tracked using near-infrared filters (excitation 769-41 nm and emission 832-37 nm). Images were analyzed and normalized using Fiji macros for picture processing and quantification expressed in fluorescence-integrated density or radiance (photons · s^-1^ · sr^-1^ · cm^-2^) integrated density. Mice were sacrificed when one limit point was reached according to our animal comity guidelines. Our comity established limit points as no more than 10% weight loss, no destress signs such as alopecia or decreasing activity, tumor size not reaching more than 1cm^3^. Tumors were considered eliminated (CE) if they were too small to be detected or harvested at the time of sacrifice. All surgical procedures and injections were performed at the CHU Sainte-Justine animal facility.

### Local irradiation in mice and immunofluorescence staining of lung tissue

Whole-thorax irradiation was performed three days after intravenous injection of 4x10^5^ non-senescent HDF. Mice were anesthetized with ketamine (100 mg/kg) and xylazine (10 mg/kg), then positioned in a lead-shielded container that exposed only the thoracic region. A single 12 Gy dose of X-rays was delivered. Ten days later, 1x10^7^ granulocytes were injected i.v. and two days later, mice were euthanized and lungs harvested for the analysis of NETs formation.

Frozen lungs were embedded in OCT compound, and 12 µm sections were prepared using a Leica cryostat. Sections were mounted on gelatin-coated slides, fixed in a 4% paraformaldehyde solution for 10 minutes and incubated overnight at 4°C with either rabbit anti-GFP polyclonal antibody (Thermo Fisher Scientific, USA, Catalog # A-11122, 1:100) to detect HDF, or goat anti-MPO polyclonal antibody (R&D Systems, USA, Catalog AF3667, 1:200), with rabbit anti-citH3 monoclonal antibody (abcam, Cambridge, UK, catalog # ab5103, 1:100) to detect NETs. Slides were washed and stained with anti-rabbit Alexa 488 and anti-goat Alexa 546 (1:500, Invitrogen), counterstained with DAPI, and mounted. Controls were stained with secondary antibodies without primary antibodies. Tissue sections were imaged using a DMR fluorescence microscope (Leica Model DMi8, Leica Microsystems CMS GmbH, Bensheim, Germany). Four fields per group were randomly selected for quantification of NETs using ImageJ.

### Mouse immune reconstitution and tumor-immune infiltrate

Human immune cells were isolated from PBMCs of healthy donors using a Ficoll-Paque (GE Healthcare) gradient and granulocytes were isolated by lysing the red blood cells pellet with 19mL of sterile deionized water for 20 seconds before adding 1 ml of sterile 20× PBS solution. Granulocytes were mixed with PBMCs at a 1:1 ratio before injection into mice. A total of 1 × 10^7^ cells (5 × 10^6^ each) were injected i.v. in 100 μl of RPMI 1640 (Wisent Bio Products). Tumors or lungs were dissociated using the human Tumor Dissociation Kit gentleMACS Octo Dissociation with Heaters (Miltenyi Biotec), according to the manufacturer’s instructions. Immune cell infiltration was determined using the following specific antibodies: mouse CD45/PE/Cy7, human CD45/BUV395, human CD3/AF700, human CD19/PE-CF594, human CD4/BB515, human CD8/BV421, human CD14/APC/H7, human CD56/BV786, and human CD127/BB700, all from BD Biosciences except human CD33/BV510 and human CD25/BV711 purchased from Biolegend (**Supplementary Table 1**). Cells were processed using an LSR Fortessa (BD Biosciences), and data analysis was performed using the FlowJo V10 (v10; Tree Star) software. Uniform manifold approximation and projection (UMAP) and clustering analyses based on flow cytometry data were performed using the licensed OMIQ online platform. The following parameters were used for the UMAP dimensional reduction: 15 neighbours, a minimum distance of 0,4, and a Euclidean metric. The meta-clustering was performed using the FLOWSOM algorithm, a Euclidean distance metric, and a k value of 7.

### NETosis assay and image analysis

To detect NETosis, we used the IncuCyte S3 (Essen Bioscience) live cell imaging system in combination with Incucyte® Cytotox red dye or green dye (Sartorius, Ann Arbor, MI, USA). Granulocytes (5 x 10^4^) were co-culture with non-senescent or senescent HDF (3 × 10^4^) in a 48-well plate. Alternatively, granulocytes were cultured in the presence of 500µL of CM collected from HDF in a 48-well plate. Incucyte® Cytotox red or green dye (250 nM) was added to the media and NETosis was monitored for 24 hours with photos taken at one-hour intervals. Cells were treated with 25 nM of PMA (Sigma Aldrich, P8139) as a positive control. NETosis was quantified by measuring the total red or green area (µm^2^ per image), and the data were analyzed using IncuCyte software. In co-culture assays, cells were incubated with or without reparixin at the specified concentration (Sigma Aldrich, SML2655).

### Tumor spheroid, invasion assay and therapy-induced senescence

To generate tumor spheroid, 5x10^3^ tumor cells were mixed with 5x10^3^ HDF (non-senescent or senescent) in a well of an ultra-low attachment 96-well plates, centrifuged for 5 minutes at 150g and then incubated at 37°C under 5% CO^2^ for six days. PBMCs with or without granulocytes (3 × 10^5^ each) were added. 24 hours later, spheroids were washed twice with PBS 1X, dissociated into single cells, and cell infiltration was determined by flow cytometric analysis using counting beads. To assess therapy-induced senescence, we first generated spheroids by seeding 1 × 10⁴ tumor cells and an equivalent number of HDFs and incubated the cells at 37 °C with 5% CO₂ for only four days to prevent necrotic core formation, as the size of spheroids increases in subsequent days of treatments. Spheroids were then treated with 0.1µM of doxorubicin or a single dose of 15 Gy IR, and six days later, granulocytes (3 x 10^5^) along with the Incucyte® Cytotox green dye were added with or without 0.8µM or reparixin. Spheroids size and NETosis were monitored for 24 hours using the IncuCyte S3 (Essen Bioscience) with images taken every hour. All images were captured at 4x magnification. Additionally, some spheroids were fixed with 4% formaldehyde and stained to detect SA-β-gal activity.

### Data collection and analysis

Single-cell RNA sequencing data were obtained from the study of Wu Y. *et al*. (20), PRJCA020880 (available at http://www.pancancer.cn/neu/). We limited our analysis on colon adenocarcinoma (COAD) and lung cancer (LC) samples before and after treatments with radiotherapy or chemotherapy. All subsequent analyses were performed using R (version 4.5.1) and RStudio (“Cucumberleaf Sunflower”). The dataset was processed with Seurat (version 5.3.0). Group comparisons were performed using Wilcoxon rank sum tests and Student’s t-tests. Gene set enrichment analysis was conducted with the clusterProfiler package (version 4.16.0) using the running-sum statistic. The KEGG database was used for pathway analysis. For data visualization, we employed the pheatmap (version 1.0.13), ggplot2 (version 4.0.0), and dittoSeq (version 1.20.0) packages in R.

Gene sets associated with NETosis were compiled from previously published studies (23–25) and analyzed in two neutrophil subpopulations, S100A12^+^(26) and MMP9^+^(27), both of which have been reported as instrumental in initiating and promoting NETosis. These genes were sorted using an unsupervised hierarchical heatmap, before and after treatment, using Ward’s method and Euclidean distance. Moreover, biplots from principal component analyses (PCA) were generated to visualize patterns of gene expression variability and to assess similarities and differences in transcriptomic profiles among samples.

### Statistical analysis

Statistical analyses were conducted using GraphPad Prism 8.0 (GraphPad Software, Inc., San Diego, CA). Data are presented as mean ± SEM. For comparisons involving three or more groups, t-tests or one-way ANOVA analysis, followed by Dunnett’s multiple comparisons test for comparisons to a control mean or Tukey’s multiple comparisons test for all pairwise comparisons. Tumor growth kinetics were analyzed using mixed-effects models, followed by Tukey’s multiple comparisons test. Statistical significance was defined as *p < 0.05, **p < 0.01, ***p < 0.001, and ****p < 0.0001.

## RESULTS

### Senescent fibroblasts interfere with the tumor immune response in humanized mice

To evaluate the impact of a senescent microenvironment on the tumor immune response, we employed a humanized mouse model in which tumor growth is delayed following the injection of autologous immune cells (20). In this model, tumor cells, HDF and immune cells were all derived from the same donor. Non-senescent or senescent HDF (previously induced by exposure to IR or RAS) were first injected intravenously through the tail vein of NSG-SGM3 mice, where they engrafted in liver and lungs **(Fig. 1A and 1B).** The following day, lung epithelial tumor cells (LEC-4T) were injected intravenously at a 4:1 ratio relative to HDF. Six days later, mice were injected intravenously with human immune cells (PBMCs and granulocytes, 5x10^6^ each). Tumor cells were engineered to express the luciferase gene, enabling tumor growth to be monitored using *in vivo* bioluminescence. Additionally, HDF were labeled with the NIR790 cytoplasmic membrane dye, facilitating their visualization and quantification **(Fig. 1B)**.

**Figure 1.**
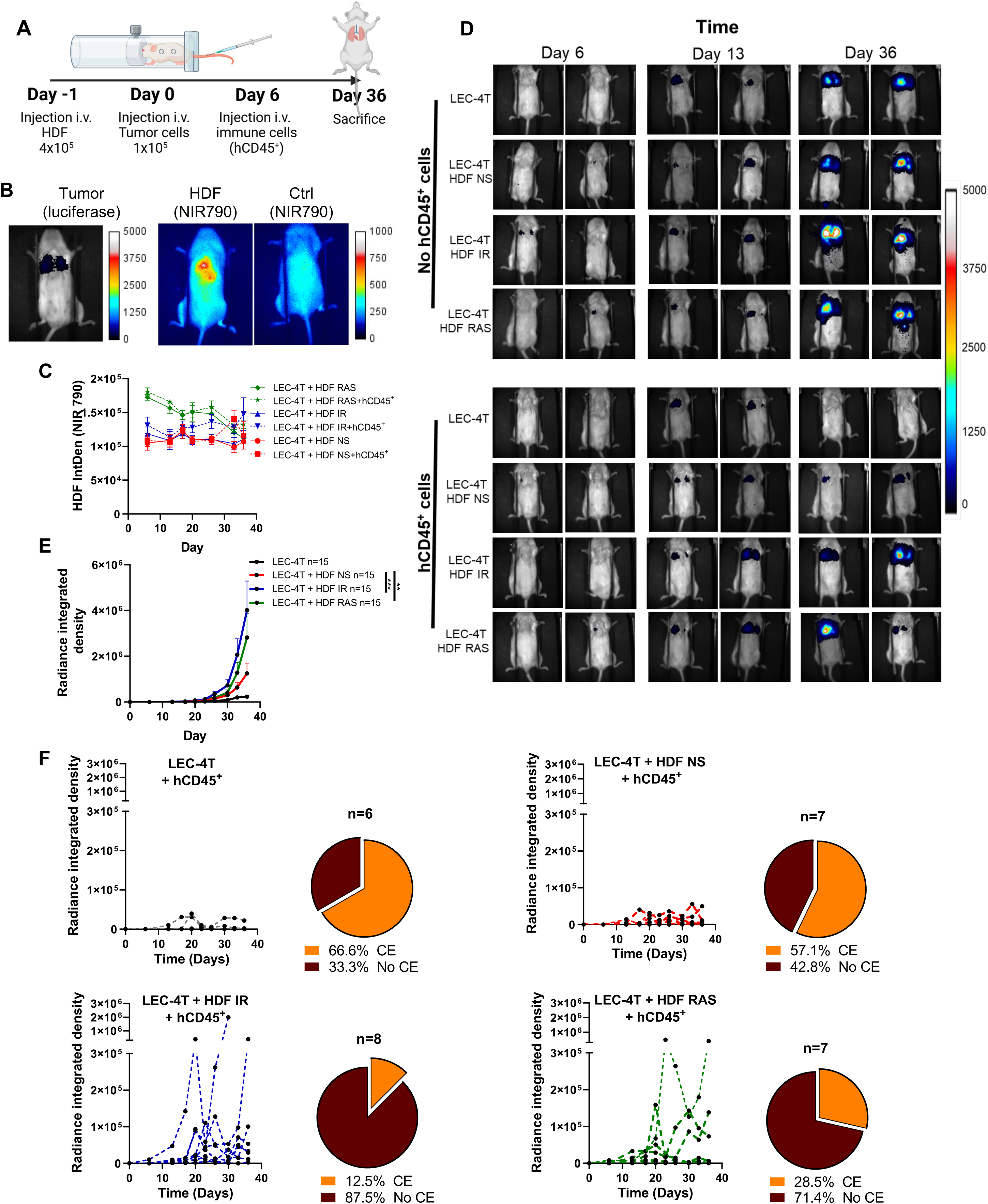
Senescent fibroblasts promote tumor growth and impair their immune rejection in lungs. **A)** Schematic illustration of the experimental workflow. NSG-SGM3 mice were injected intravenously (i.v) with 4x10^5^ non-senescent or senescent HDF. The next day, 1x10^5^ tumor cells (LEC-4T) were injected i.v.. Six days after tumor cell injection, mice were injected i.v with hCD45^+^ cells (5x10^6^ PBMCs and 5x10^6^ granulocytes each/mouse). Tumor growth was evaluated weekly until day 36, when mice were sacrificed and their lungs collected for analysis. **B)** Representative *in vivo* bioluminescence images of mice injected with luciferase-expressing LEC-4T and HDF stained with NIR790 dye. Also shown is a mouse without HDF. **C)** Graph showing the integrated density of HDF stained with NIR790 and tracked over time in LEC-4T tumors growing in NSG-SGM3 mice injected (dashed line) or not injected (solid line) with hCD45^+^ cells. Non-senescent (NS) HDF are shown in red, HDF exposed to IR in blue, and HDF expressing RAS in green. Data are presented as mean ± SEM from 8–9 tumors. **D)** Representative bioluminescence images of tumor-bearing mice with or without hCD45^+^ cells. Mice were injected with LEC-4T alone or with non-senescent or senescent HDF (IR or RAS). Images were captured on days 6, 13, and 36. **E)** Growth curves of LEC-4T tumors injected alone (black) or with non-senescent HDF (red), senescent HDF induced by irradiation (blue), or senescent HDF induced by RAS (green) in mice without hCD45^+^ cells. Each line represents the mean tumor growth ± SEM for 15 mice in each group over 36 days. Statistical analyses were performed using a mixed-effects model followed by Tukey’s multiple comparison test. **p < 0.01; ***P < 0.001. **F)** Tumor growth curves for individual mice injected with hCD45^+^ cells. LEC-4T tumor cells injected alone are shown in black (n = 6), with non-senescent HDF in red (n = 7), with senescent HDF induced by irradiation in blue (n = 8), or with senescent HDF induced by RAS in green (n = 7). Also shown is a pie chart illustrating the mean percentage of mice that achieved complete tumor elimination (CE) after immune reconstitution.

Engraftment of HDF remained stable over a five-week period following the injection of immune cells, indicating that most HDF persist and are not eliminated by immune cells **(Fig. 1C)**. As expected, we observed that HDF, particularly senescent HDF, stimulated tumor growth compared to mice injected with LEC-4T tumor cells alone **(Fig. 1D and 1E)**. However, the injection of immune cells delayed or completely eliminated tumors (CE) in 66.6% and 57.1% of mice injected with tumor cells alone or with non-senescent HDF, respectively **(Fig. 1F).** In contrast, complete tumor elimination was limited to 12.5% and 28.6% in mice that received senescent HDF induced by IR or RAS respectively **(Fig. 1F).** The negative impact of senescent HDF on the tumor immune response was not restricted to LEC-4T tumors, as we observed similar results using the A549 lung cancer cell line and allogenic immune cells **(Supplementary Fig. 1).** These findings indicate that the presence of senescent HDF not only enhances tumor growth, but it also impairs tumor rejection in humanized mice.

### Senescent fibroblasts modify the tumor immune infiltrate

To determine whether impaired tumor rejection is associated with reduced immune cell infiltration, we collected, weighted, and dissociated tumor-bearing lungs to quantify the infiltration of hCD45^+^ immune cells at the time of sacrifice. Cell counts obtained by flow cytometry and normalized to the lung weight showed that mice injected with senescent HDF tended to have a decreased infiltration of hCD45^+^ **(Fig. 2A).** However, the proportion of infiltrated hCD33^+^ myeloid cells was significantly increased in mice injected with senescent HDF, showing a 2.7-fold increase with HDF induced by IR and a 2-fold increase with HDF induced by RAS, respectively (**Fig. 2B-D).** Such an increase was associated with a concomitant decrease in other immune cell subsets, such as CD4^+^ T cells and NK cells **(Fig. 2B and 2C)**.

**Figure 2.**
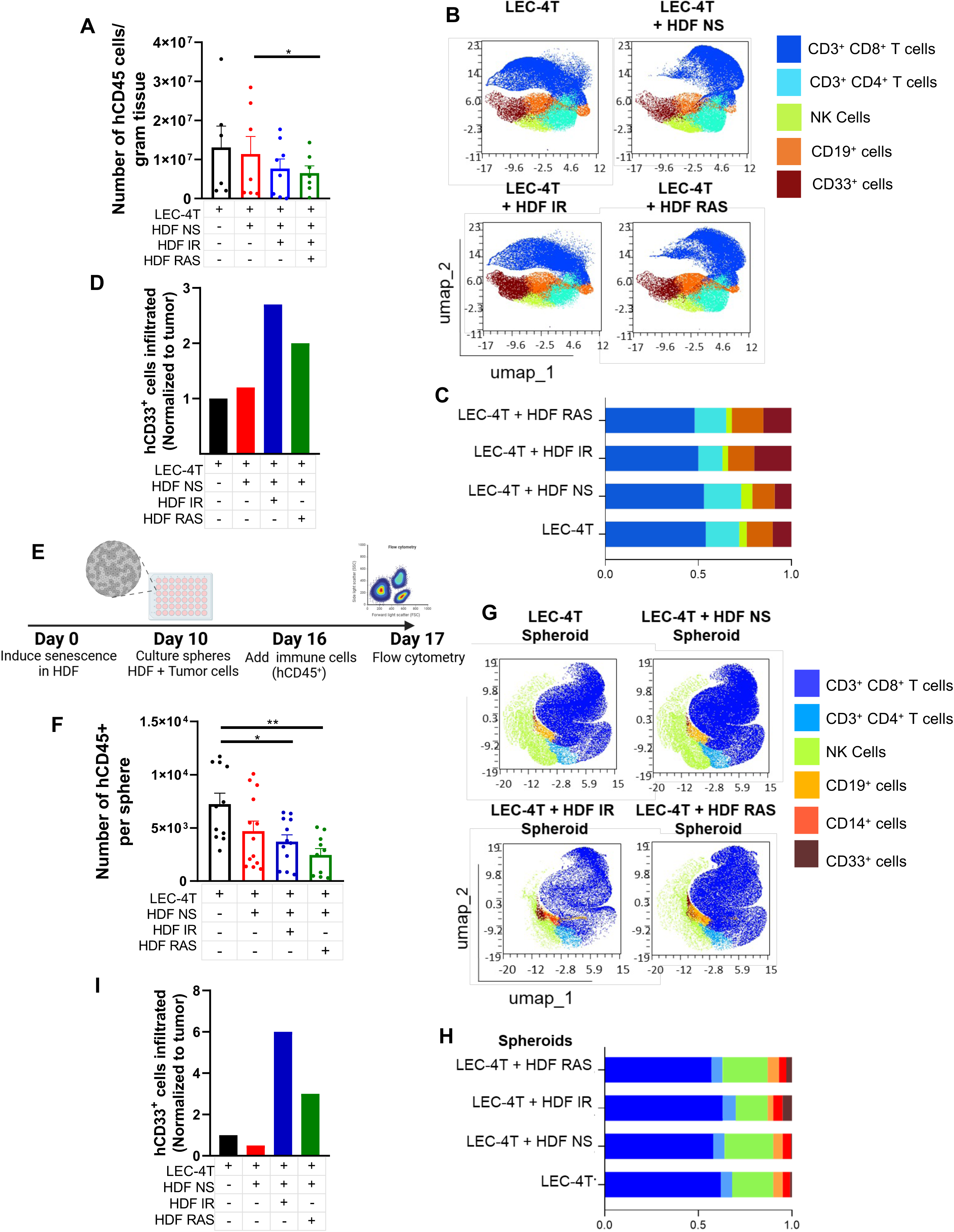
Senescent fibroblasts increase the recruitment of myeloid cells in the lung tumor microenvironment and tumor spheroids. **A)** Graph depicting absolute counts of lung-infiltrating hCD45^+^ cells per gram of tissue, determined by flow cytometry for the experiment shown in Figure 1. Each dot indicates the count for an individual mouse. Data are presented as mean ± SEM. Statistical significance was assessed using one-way ANOVA followed by Dunnett’s multiple comparisons test. *p < 0.05. **B)** Two-dimensional UMAP plot illustrating merged clustering of lung-infiltrating subtypes of hCD45^+^ cells in mice bearing LEC-4T alone or with HDF (non-senescent and senescent). **C)** Histograms showing the frequencies of cell composition from the clustering depicted in panel B. **D)** Graph showing the proportion of hCD33^+^ cells infiltrating the lungs. Data were normalized to the condition with LEC-4T tumor cells injected without HDF. **E)** Schematic representation of the tumor spheroid model. HDF were induced to senescence by exposure to ionizing radiation (IR) or activation of the RAS oncogene. After 10 days, tumor cells (LEC-4T) alone, or an equal number of LEC-4T and either non-senescent or senescent HDF (5,000 cells each), were used to generate monospheroids or mixed-cell spheroids. The spheroids were allowed to form and mature for 6 days, followed by co-culture with 3 x 10⁵ hCD45^+^ cells for 24 hours. Spheroids were then washed prior to be dissociated and immune cell infiltration was assessed by flow cytometry. **F)** Graph showing the number of hCD45^+^ cells infiltrating spheroids from each group after dissociation. Each data point represents an individual spheroid. Statistical comparisons among groups were conducted using one-way ANOVA with Tukey’s multiple comparisons test. *p < 0.05 **p < 0.01. **G)** Two-dimensional UMAP visualization illustrating consolidated clustering of spheroid-infiltrating subtypes of hCD45^+^ cells across groups. **H)** Histograms illustrate the frequency distributions of immune cell compositions resulting from the clustering analysis presented in panel **G**. **I)** Graph displaying the proportion of hCD33^+^ cells infiltrating spheroids. Data were normalized to spheroids generated with LEC-4T alone.

Given the differences in residual tumor sizes at the time of sacrifice, we sought to assess immune cell infiltration using a more defined model. To this end, we established a *in vitro* tumor spheroid model in which LEC-4T tumor cells are co-cultured with non-senescent or senescent HDF in round-bottom 96-well plates. After six days, once spheroids had formed, hCD45^+^ immune cells (PBMCs and granulocytes) were added and allowed to infiltrate for 24 hours prior to the dissociation of the spheroid and the quantification of immune cell using flow cytometry **(Fig. 2E)**. Consistent with the observations in lungs, we detected a reduction in the overall infiltration of hCD45^+^ immune cells in spheroids containing senescent HDF **(Fig. 2F)**. Furthermore, spheroids containing senescent HDF showed increased infiltration of hCD33^+^ myeloid cells, particularly in those containing IR-induced senescent HDF **(Fig. 2G-I)**.

In addition, we evaluated immune cell infiltration 24 hours after the injection of immune cells in mice bearing pre-established lung metastasis **(Supplementary Fig. 2A)**. Similarly to what we observed with spheroids, there was a higher proportion of hCD33^+^ infiltrating myeloid cells in tumors grown in the presence of senescent HDF compared with non-senescent HDF **(Supplementary Fig. 2B and 2C)**. By measuring immune cell infiltration shortly after their injection (24 hours) in both mice and tumor spheroids, it allowed us to measure the impact of senescence on cell recruitment independently of cell proliferation if any. Overall, these results suggest that a senescent tumor microenvironment favors the recruitment of myeloid cells while impairing whole immune cell recruitment.

### Senescent fibroblasts induce NETosis through their SASP

Our observation that more myeloid cells are recruited despite an overall reduction in the tumor immune infiltrate led us to hypothesis that senescent HDF may trigger the formation of NETs. To test this hypothesis, we co-cultured human granulocytes with non-senescent or senescent HDF at a 2:1 ratio in the presence of a fluorescent cell impermeant DNA-binding reagent (Incucyte® Cytotox Red) to quantify NETosis using live imaging. We found that co-cultures with senescent HDF, regardless of the inducer, stimulated NETs formation compared to conditions with non-senescent HDF or granulocytes alone **(Fig. 3A and 3B)**.

**Figure 3.**
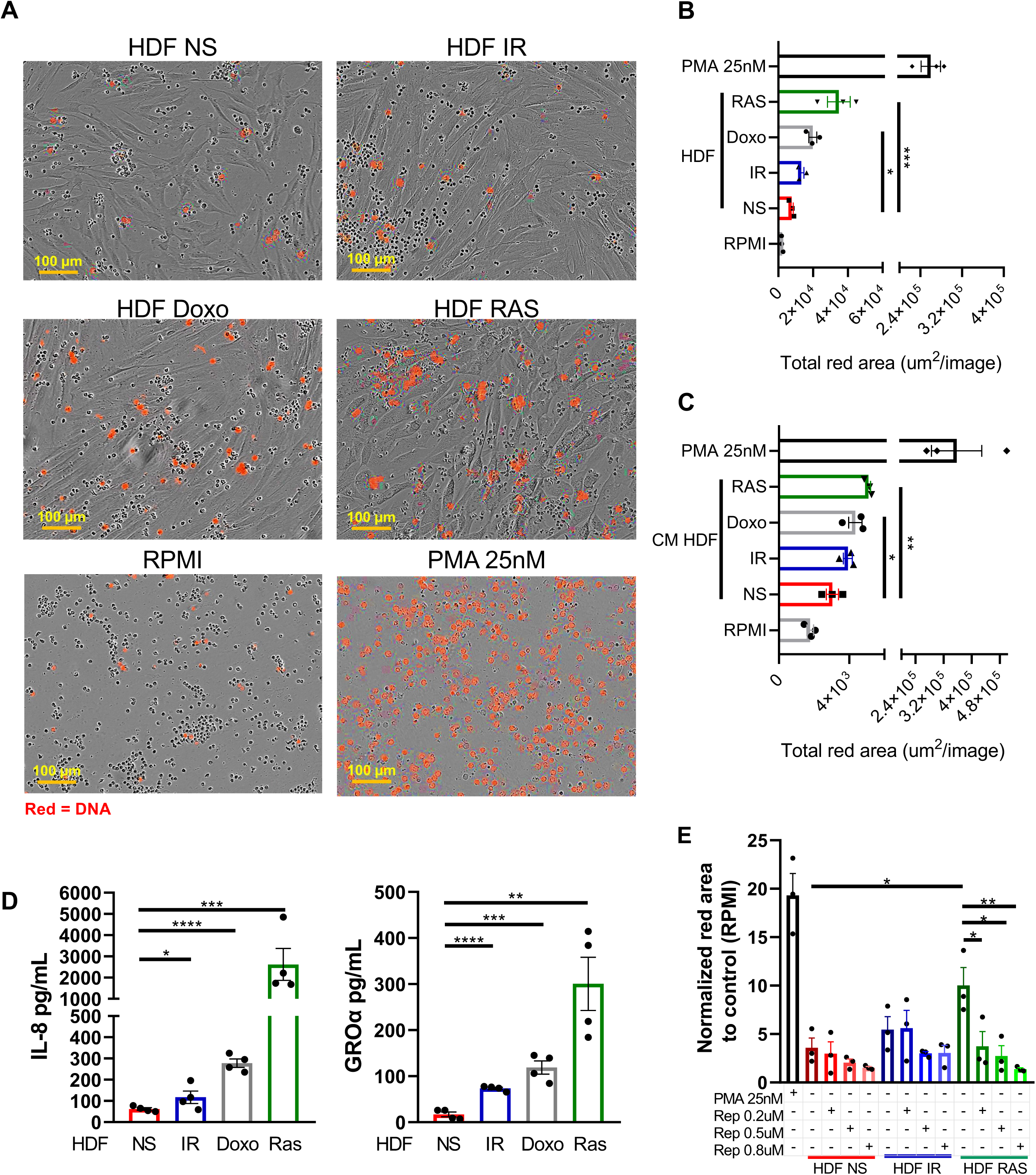
Senescent fibroblasts induce NETosis through their SASP. **A)** Representative images captured by the IncuCyte using a 10X objective showing granulocytes cultured alone (RPMI), stimulated with PMA (25 nM), or co-cultured with HDF (non-senescent or senescent) at 5 hours. Neutrophils undergoing NETosis are visualized as red clouds of extracellular DNA that fluoresce in the presence of the Cytotox Red Reagent. The scale bar represents 100 µm. **B)** Graph showing the total red fluorescence area observed in each group shown in panel A. The images were analyzed using the IncuCyte Live-Cell Analysis System. Data are presented as the mean ± SEM from three independent biological replicates. A one-way ANOVA, followed by Dunnett’s multiple comparisons test, was used to determine statistical significance. *p < 0.05; ***p < 0.001. **C)** Graph representing the total red fluorescence area after culturing granulocytes alone (RPMI), stimulated with PMA (25 nM), or using the conditioned media (CM) of non-senescent or senescent HDF with Cytotox Red Reagent to detect extracellular DNA at 7 hours. Data are presented as the mean ± SEM from three independent biological replicates. A one-way ANOVA, followed by Dunnett’s multiple comparisons test, was used to determine statistical significance. *p < 0.05; **p < 0.01. **D)** Concentration of IL-8 and Gro-α (pg/mL) from the serum-free CM of non-senescent or senescent HDF, normalized to the number of cells used. Data are presented as the mean ± SEM from four independent biological replicates. A one-way ANOVA, followed by Dunnett’s multiple comparisons test, was used to determine statistical significance. *p < 0.05; **p < 0.01; ***p < 0.001; ****p < 0.0001. **E)** Quantification of NETs using the IncuCyte Live-Cell Analysis System and Cytotox Red Reagent. Granulocytes were stimulated with PMA (25 nM) or co-cultured with HDF (non-senescent or senescent). Three different concentrations of reparixin (0.2 µM, 0.5 µM, and 0.8 µM) were added to the cultures. Results were normalized to the red fluorescence area of granulocytes cultured alone (RPMI) and are presented as the mean ± SEM from three independent biological replicates. A one-way ANOVA, followed by Tukey’s multiple comparisons test, was used to determine statistical significance. *p < 0.05; **p < 0.01.

We next sought to determine whether NETs release was attributable to the SASP of HDF. To test this possibility, we measured NETs formation in presence of conditioned media (CM) collected from non-senescent or senescent HDF. Our results revealed that CM collected from senescent HDF is sufficient to enhance NETs formation **(Fig. 3C)**. These findings may be explained by the presence of CXCL8 (IL-8) and CXCL1 (Gro-α), two chemokines often observed in the SASP of senescent HDF and known to induce NETosis (11). Indeed, we observed that the concentration of both chemokines was increased in the CM of senescent HDF, particularly in cells induced to senesce by RAS **(Fig. 3D)**. Given that CXCL1 and CXCL8 signal through CXCR1 and CXCR2 on neutrophils, we next evaluated if we could block the formation of NETs in presence of reparixin, a specific allosteric inhibitor of CXCR1 and CXCR2 (28). We found that reparixin, in a dose-dependent manner, was able to completely abolish the formation of NETs **(Fig. 3E)**. Altogether, these results indicate that senescent HDF, regardless of the inducer, enhances NETs formation through their SASP.

### NETosis triggered by senescent fibroblasts reduces the infiltration of immune cells in tumor spheroids and in tumor-bearing mice

We next sought to investigate whether senescence-induced NETs are responsible for the overall decrease in immune cell infiltration observed in our tumor models. To address this question, we first used tumor spheroids assembled with non-senescent or senescent HDF and determined if the infiltration of immune cells was inversely correlated with the release of NETs **(Fig. 4A)**. In this model, HDF are predominantly located at the core regions of the spheroid **(Fig. 4B)**. As expected, more extracellular DNA was observed surrounding spheroids containing senescent HDF, compared to non-senescent HDF, an effect inhibited by the addition of reparixin **(Fig 4C and 4D)**.

**Figure 4.**
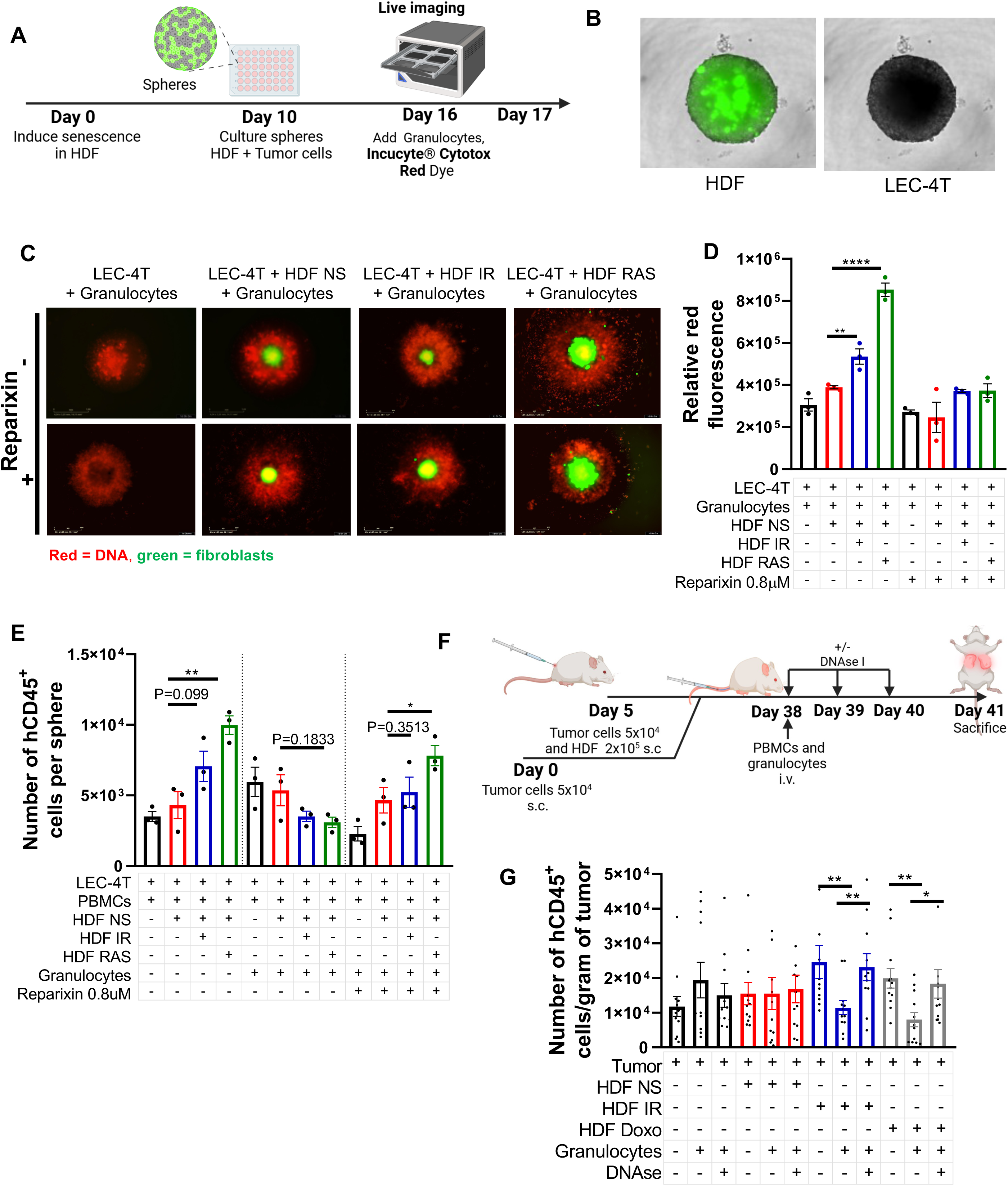
Senescence-induced NETosis decreases the infiltration of immune cells in tumor spheroids and tumor-bearing mice. **A)** Schematic representation of LEC-4T tumor spheroids assembled as described in figure 2E. In brief, 4 × 10^5^ granulocytes were added to mature spheroids along with the Cytotox Red Reagent dye and NETosis around spheroids was monitored using the IncuCyte Live-Cell System, with scans taken at 2-hour intervals for up to 24 hours. **B)** Representative images (at 4X magnification) of spheroids on day 16 before adding granulocytes. HDF (expressing EGFP) are shown in green. **C)** Representative images (at 4X magnification) showing NETosis (in red) and HDF bearing spheroids (in green) 24 hours after the addition of granulocytes. Where indicated, Reparixin (0.8 µM) was added to reduce NETs formation. **D)** Graph showing the red fluorescence intensity for each group in panel **C**. Each dot represents an independent experiment and reflects the mean value calculated from 12 spheres per condition. Statistical analysis between groups was performed using one-way ANOVA with Dunnett’s multiple comparisons test. **p < 0.01; ****p < 0.0001. **E)** Spheroids were co-cultured with PBMCs with or without granulocytes and reparixin (0.8 µM). The graph shows the number of hCD45⁺ cells infiltrating spheroids in each indicated group after dissociation and flow cytometric analysis. Data are presented as the mean ± SEM from three independent biological replicates and reflect the mean value calculated from 8 spheres per condition. Statistical analysis between groups was performed using one-way ANOVA with Tukey’s multiple comparisons test. *p < 0.05; **p < 0.01. **F)** Schematic of the *in vivo* experimental design. Two groups of NSG-SGM3 mice were injected subcutaneously with either 5 × 10^4^ tumor cells (LEC-4T) alone or five days later with tumor cells mixed with non-senescent or senescent HDF (2 × 10^5^ cells). Mice were injected with tumor cells alone to achieve comparable tumor volumes across all experimental groups prior to the injection of immune cells. Tumor growth was evaluated weekly until day 38, at which point mice were intravenously injected with PBMCs (5 × 10^6^ cells/mouse) and were indicated granulocytes (5 × 10^6^ cells/mouse). On day 38 and for the subsequent two days, mice were treated with DNase I (2.5 mg/kg). On day 41, mice were sacrificed and tumors were collected for analysis. **G)** Graph showing the number of tumor-infiltrating hCD45⁺ cells per gram of tumor. Each dot corresponds to counts in an individual tumor. Statistical analysis between groups was performed using one-way ANOVA with Tukey’s multiple comparisons test. *p < 0.05 **p < 0.01.

To determine whether these NETs prevent immune cell infiltration, we added PBMCs with or without granulocytes to the spheroids and quantified the number of infiltrating hCD45^+^ immune cells 24 hours later. Our results clearly demonstrate that senescent induced-NETs inhibit the infiltration of hCD45^+^ immune cells, an effect that was completely reversed by the addition of reparixin **(Fig. 4E)**. Further examination of the immune cell subsets infiltrating the spheroids, revealed that NK cells and CD33^+^ myeloid cells were the most affected **(Supplementary Fig. 3)**. These results confirm the recruitment of myeloid cells by senescent HDF and establish a link between NETs induction and an overall impairment of immune cell recruitment.

To validate these results, we co-injected LEC-4T tumor cells with non-senescent or senescent HDF subcutaneously into the flanks of NSG-SGM3 mice. Tumor cells alone were injected 5 days earlier than those co-injected with HDF to compensate for their slower growth. Tumors were allowed to grow for up to 38 days, reaching an average size of 400 mm^3^. Mice were then injected intravenously with PBMCs, either alone or with granulocytes. To inhibit NETs formation, a subset of mice also received intramuscular injections of DNase I for 3 days. At the time of sacrifice, immune cell infiltration was analyzed in dissociated tumors **(Fig. 4F)**. Our results revealed a decrease in hCD45^+^ cell infiltration in tumors containing senescent HDF when granulocytes were injected, consistent with the results obtained in spheroids **(Fig. 4G)**. Such a decrease was partially reversed in mice treated with DNase I **(Fig. 4G)**.

### Therapy-induced senescence increases NETosis and impairs immune cell infiltration

We next sought to determine whether therapy-induced senescence (TIS) in lungs, a model that replicates cancer treatments, would lead to NETosis in vivo. To do so, we first injected 4×10^5^ HDF intravenously into NSG-SGM3 mice, knowing that the cells would engraft mostly into the lungs (**Fig. 1B**). Three days later, lungs were locally irradiated to induce senescence. We waited ten days for the senescence phenotype to develop and then injected 1×10^7^ granulocytes intravenously **(Fig. 5A)**. Two days later, we looked for evidences of NETosis in lungs sections by immunofluorescence stainings for human myeloperoxidase (MPO) and human citrullinated histone H3 (citH3), two specific markers for neutrophils and NETosis **(Fig. 5B)**. We found that mice injected with HDF exhibited increased NETosis compared to those without HDF, even in absence of irradiation (**Fig. 5C**). However, more NETosis was observed in lungs containing HDF and exposed to irradiation as quantified from MPO and citH3 fluorescent signals **(Fig. 5C).** Of note, injection of granulocytes in absence of HDF only lead to a modest increase in NETosis even after exposure to irradiation, suggesting the SASP from mouse tissues does not trigger NETosis of human granulocytes.

**Figure 5.**
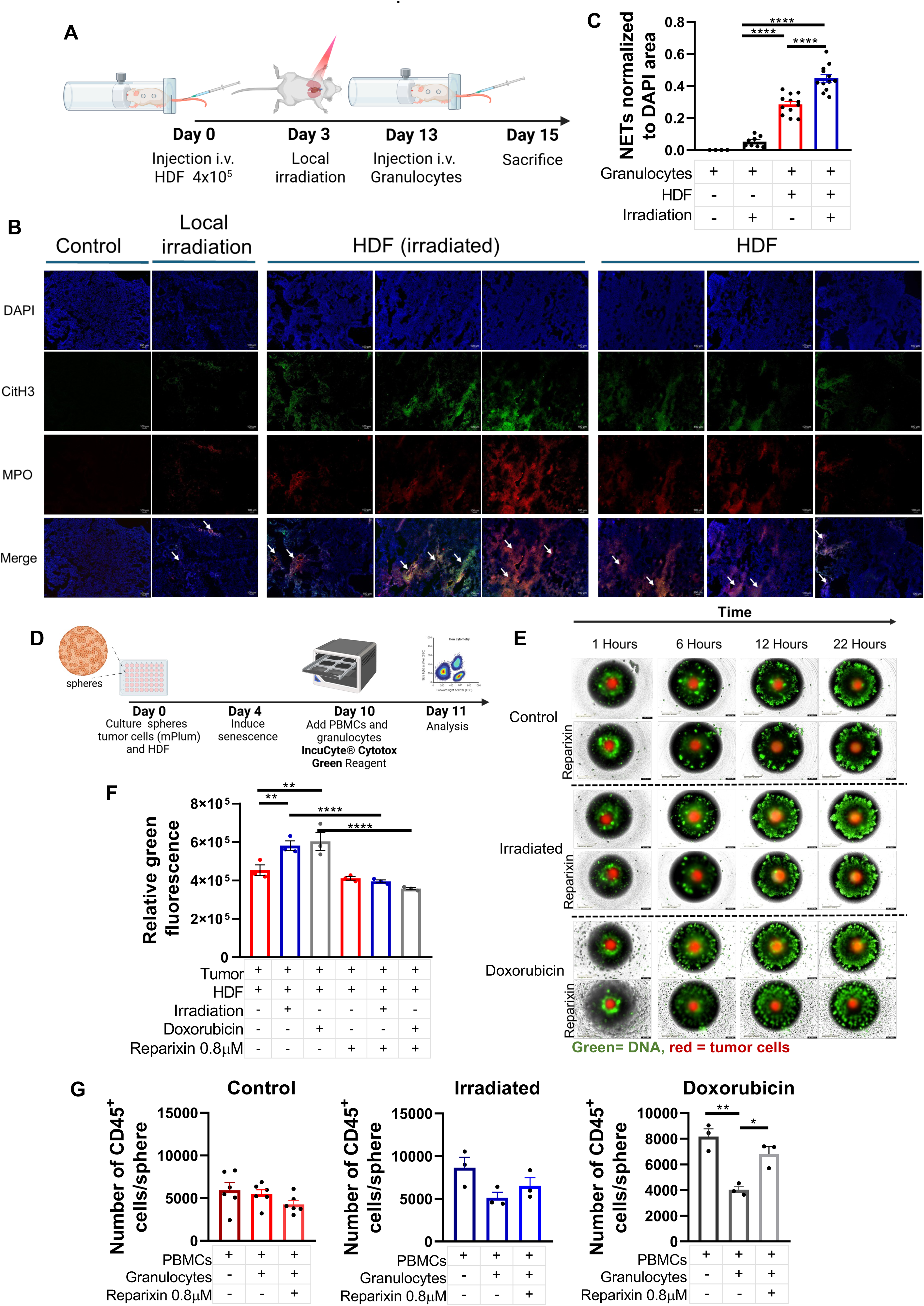
Therapy-induce senescence increases NETosis and impairs immune cell infiltration in tumor spheroids. **A)** Schematic representation of the *in vivo* experimental design. NSG-SGM3 mice were injected intravenously (i.v.) with 4×10^5^ non-senescent HDF. Three days later, mice received a single dose of whole-thorax radiation (12 Gy). Ten days later, mice were injected i.v. with 1x10^7^ granulocytes and 48 hours later mice were sacrificed and lungs collected for analysis. **B)** Representative immunofluorescence staining of NETs formation (arrow) formed by co-localization of citrullinated Histone 3 (CitH3 in green), myloperoxidase (MPO in red) and DNA (DAPI in blue) from irradiated and non-irradiated lung tissues previously injected or not with HDF. The scale bar represents 100 μm. **C)** Quantification of NETs formation detected by the fluorescence signals for MPO and CitH3 colocalization. Data were normalized by the total area occupied by the DAPI stain in that same region. Statistical analysis between groups was performed using one-way ANOVA with Tukey’s multiple comparisons test. ****P < 0.001. **D)** Schematic representation of the tumor spheroid model. In brief, LEC-4T tumor cells expressing mPlum were mixed with an equal number of HDF (1x10^4^ cells each) to form mixed-cell spheroids. Four days later, spheroids were treated with doxorubicin (0.1µM) or irradiated (15Gy) to induce senescence. Six days after treatment, PBMCs (and granulocytes (3X10^5^ cells each) were added along with the Cytotox Green Reagent. Where indicated spheroids were treated with reparixin (0.8 µM) for 24 hours and immune cell infiltration within the spheroids was quantified by flow cytometry. **E)** Representative images (at 4X magnification) of mixed-cell spheroids showing tumor cells in red and NETs visualized as fluorescent extracellular DNA in green. Reparixin (0.8 µM) was added to reduce NETs formation. The scale bar represents 100 μm. **F)** Graph showing the green fluorescence intensity from each group shown in panel E. Each dot represents an individual experiment, reflecting the mean value calculated from three spheres per condition. Statistical analysis between groups was performed using one-way ANOVA with Tukey’s multiple comparisons test. **p < 0.01; ****P < 0.001. **G)** Graphs depicting the number of hCD45⁺ cells infiltrating spheroids from each indicated group. Each dot represents an independent experiment and reflects the mean value calculated from 8 spheres per condition. Statistical analysis between groups was performed using one-way ANOVA with Tukey’s multiple comparisons test. *p < 0.05; **p < 0.01.

To determine whether NETosis also impaired immune cell infiltration in the context of TIS, we exposed pre-formed spheroids, composed of LEC-4T tumor cells and non-senescent HDF, to either IR or doxorubicin. Treated spheroids were incubated for 6 days to allow the senescence phenotype to develop, after which PBMCs and granulocytes were added to the culture with or without reparixin **(Fig. 5D)**. Successful induction of senescence was confirmed by detecting SA-β-gal activity in the spheroid and by tracking cell growth using live imaging **(Supplementary Fig. 4)**. Increased NETs formation was observed at the periphery of spheroids following treatments, an effect that was reversed by reparixin **(Fig. 5E and 5F)**. Dissociation of spheroids and quantification of immune cells revealed impaired infiltration of hCD45^+^ immune cells in the presence of granulocytes, a defect that was mitigated by reparixin **(Fig. 5G)**. These results demonstrate that TIS promotes NETosis and reduces immune cell infiltration.

To further support these observations, we next analyzed published single-cell RNA sequencing dataset of patient tumor neutrophils (21). From this analysis, we selected colon adenocarcinoma and lung cancer samples with pre- and post-genotoxic treatment data. Gene set enrichment analysis (GSEA) identified significant upregulation of pathways related to neutrophil chemotaxis, activation, and migration, associated with NETosis, in samples collected from patients post-treatments **(Supplementary Fig. 5A and 5B)**. These findings are consistent with the increased expression of 25 key NETosis-related genes. For examples, PADI4 (Peptidyl Arginine Deiminase 4), an enzyme essential for citrullination (29, 30), and PTAFR (Platelet-Activating Factor Receptor), are implicated in driving the release of NETs in the TME (31, 32) **(Supplementary Fig. 5C and 5D)**. The expression patterns of these genes are presented in a heatmap across two neutrophil subpopulations, S100A12+ and MMP9+, each previously linked to NETosis initiation and promotion (25, 26). Correspondingly, the PCA biplot indicates a pronounced distinction in gene expression between the two subpopulations before and after treatment **(Supplementary Fig. 5C and 5D)**. These results support the idea that NETosis is increased after TIS.

### Senescence-induced NETosis promotes metastasis in lungs

Senescence and NETosis have been reported to promote tumor metastasis independently (33, 34). Whether NETosis is a mechanism by which senescent cells increase metastasis formation is unknown. To investigate this, we injected non-senescent or IR-induced senescent HDF into the tail vein of NSG-SGM3 mice. Five and seven days later, we injected granulocytes and LEC-4T tumor cells respectively, with tumor cells expressing the firefly luciferase and mPlum genes. Starting the following day, subgroups of mice received intramuscular injections of DNAse I to reduce NETs formation **(Fig. 6A)**. Using this protocol, we expected that HDF would predominantly be trapped in the lungs (as shown in **Fig. 1B**), and that NETs formed in the presence of senescent HDF would lead to increased metastasis.

**Figure 6.**
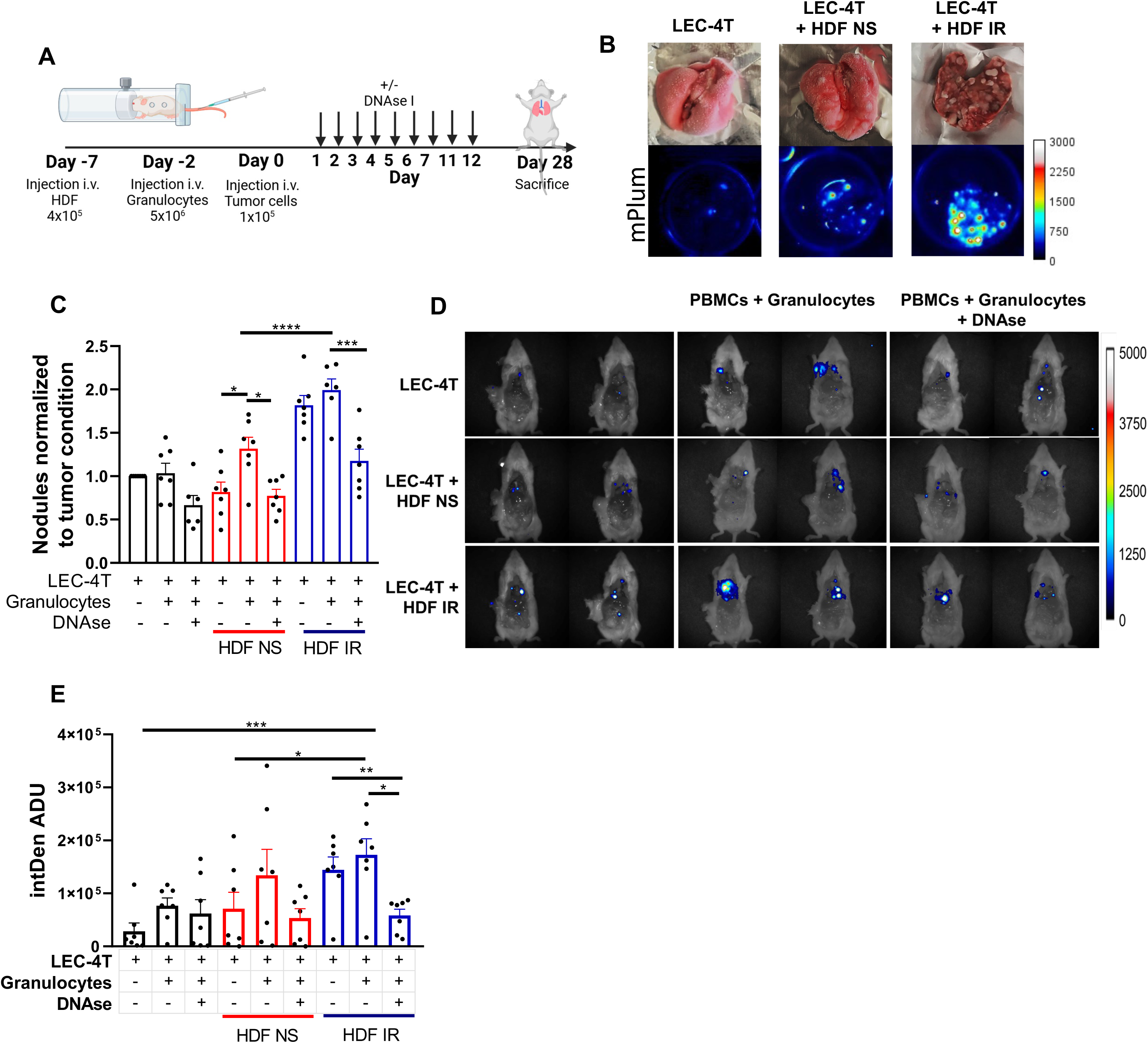
NETosis induced by senescent fibroblasts enhances lung metastasis. **A)** Schematic illustration of the experimental design. In brief, NSG-SGM3 mice were injected intravenously (i.v.) with 4 x10^5^ non-senescent or senescent HDF (IR), five days later with 5 x10^6^ granulocytes and two days later (day 7) with 1 x10^5^ LEC-4T tumor cells. Mice were subsequently treated with DNAse I (2.5 mg/kg) for twelve days. Tumor growth was assessed *in vivo* via bioluminescent imaging every week until day 28, when mice were sacrificed and lungs collected for analysis. **B)** Representative photographs of dissected lung tissue from each group of mice, showing metastatic surface nodules. Fluorescent imaging of the lungs depicting tumors (expressing mPlum) is also shown to assess metastases located deeper within the lung parenchyma. **C)** Graph presenting the number of surface nodules per lung in each group normalized to the number of nodules observed when tumor cells were injected alone. Each dot corresponds to an individual lung collected. Statistical significance was determined using a one-way ANOVA followed by Tukey’s multiple comparisons test. *p < 0.05; ***P < 0.001; ****P < 0.0001. **D)** Representative *in vivo* bioluminescence images of mice captured after the body cavities were opened and the lungs removed. **E)** Graph showing the quantified bioluminescence signal intensity from images depicted in panel **D**. Each dot corresponds to an individual mouse. Statistical significance was determined using a one-way ANOVA followed by Tukey’s multiple comparisons test. *p < 0.05; **p < 0.01; ***P < 0.001.

Our results showed that at the time of sacrifice, twice as many metastatic nodules were macroscopically visible in the lungs of mice injected with senescent HDF compared to those injected with non-senescent HDF **(Fig. 6B and C)**. Conversely, mice treated with DNAse I showed a marked reduction in the number of metastasis **(Fig. 6C).** Metastasis formation was also increased outside of the lungs, primarily in the thoracic cavity, as determined by bioluminescence signal after the removal of lungs (**Fig. 6D)**. As observed in lungs, quantification of whole-body bioluminescence signal intensity was also increased in mice injected with senescent HDF and granulocytes **(Fig. 6E)**. Again, metastasis burden was decreased in mice injected with DNAse I **(Figure 6E).** These data demonstrate that senescence-induced NETs is a mechanism that contributes to increased metastasis formation.

## DISCUSSION

In this study, we uncover an unexpected deleterious consequence of the senescence phenotype in the context of the tumor immune response. Using a humanized orthotopic lung tumor model and autologous immune cells, we found that senescent HDF impair the tumor immune response and promote metastasis formation. Mechanistically, we demonstrated that senescent HDF, through their SASP, recruit myeloid cells, induce the formation of NETs and negatively impact immune cell infiltration into tumors *in vivo* and spheroids *in vitro*.

Senescent cells have been reported to attract myeloid cells, such as macrophages, neutrophils, and mast cells, thereby facilitating tumor proliferation by establishing an immunosuppressive microenvironment (14, 35, 36). Additionally, it has been recently shown that senescent fibroblasts, in the context of TIS, can awakens dormant cancer cells and promote metastatic relapse through the induction of NETs (37). In this study, using adoptive transfer of human granulocytes, we demonstrated that senescent HDF recruit CD33^+^ myeloid cells and induce NETosis. We identified two important SASP factors, CXCL8 (interleukin-8) and CXCL1 (GRO-α), that by binding to CXCR1 and CXCR2 play an important role in the formation of NETs (17). Indeed, we were able to inhibit NETosis using reparixin, a non-competitive allosteric inhibitor targeting the CXCR1 and CXCR2 receptors. Intriguingly, while NETs formation is deleterious in the context of the tumor immune response, it has been shown to be beneficial in the context of retinopathy, highlighting the pleiotropic and tissue-dependent effects of cellular senescence (38).

The impact of the SASP on the tumor immune response is complex and highly context-dependent. This is best illustrated by the work of Ruscetti et al., who showed that therapy-induced senescence, using the same drug combination, leads to the recruitment of distinct immune cells in different tumor models, NK cells in lung tumors and T cells in pancreatic tumors (39, 40). In our model, we observed that the infiltration of NK cells was significantly inhibited by the generation of NETs, an effect that was partially reverse by reparixin. Therefore, it is possible that the impact of senescence-induced NETs may have variable consequences depending on the tumor model used. Intriguingly, we observed that senescent HDF led to increased metastasis even in the absence of human granulocyte and that such an increase was reversed by the injection of DNAse I **(Fig. 6E)**. This suggests that the SASP of senescent HDF may be capable of inducing NETs from mouse neutrophils, although this remains to be determined.

One limitation of our study is the we could not evaluate the impact of TIS-induced NETs in tumor growth in mice, due to the challenges in synchronizing tumor growth and regression after treatment in lungs. We tried repeating the experiment using subcutaneous tumors where it would be easier to assess growth after therapy, however this was not possible given the weak immune response against subcutaneous tumors. Still, we identified an increase in NETosis markers after local irradiation in mice. Additionally, by analyzing a published single-cell RNA-seq dataset of neutrophils from patient tumors (21), we identified a correlation between treatment and upregulation of NETosis-related genes.

In conclusion, our results demonstrate that TIS, regardless of the inducer, negatively impacts the tumor immune response in humanized lung cancer models. These findings also provide insights into how alterations in the tumor microenvironment in response to therapy may adversely impact cancer treatment outcomes. Collectively, the data support the one-two punch strategy, which involves eliminating TIS cells through senolytic approaches. By indirectly inhibiting NETs, this strategy may act synergistically with existing cancer immunotherapies (14, 41, 42).

## Acknowledgments

We are grateful to the flow cytometry and animal facility for providing technical support and Dr Roni F. Rayes for technical advises. We also thank le réseau ThéCell from the Fonds de la recherche du Québec – Santé (FRQS) for supporting the iPSC platform.

## Author contributions

M.C.B. and G.M.N. performed experiments. M.C.B., A.S and C.C.G. analyzed the data. M.C. and C.B. designed the studies and analyzed the data. G.F. edited the manuscript and analyzed the data. M.C.B. and C.B. wrote the manuscript.

---

**Supplementary Figure 1.**
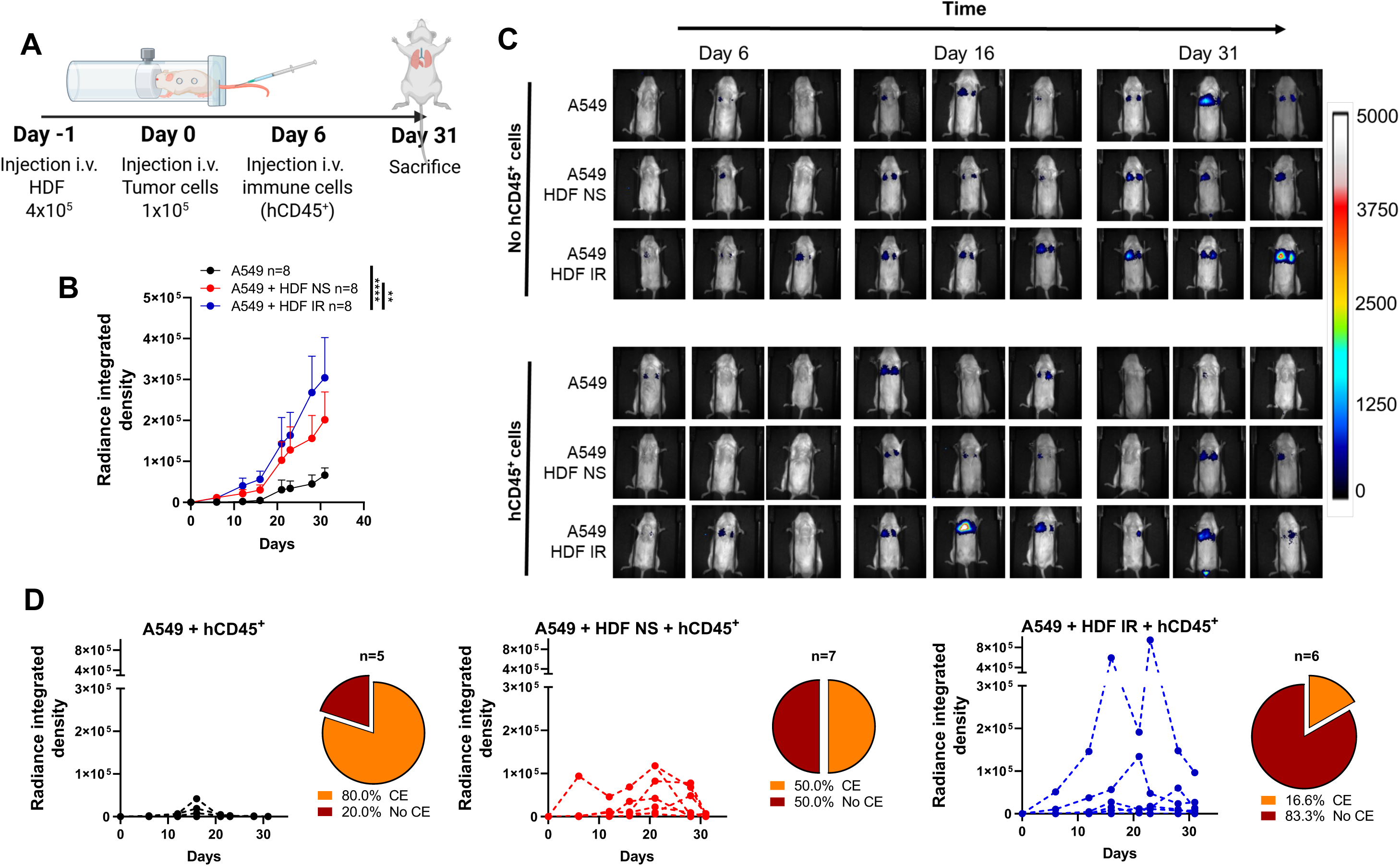
Senescent fibroblasts promote the growth of A549 tumors and impair their immune rejection in lungs. A) Schematic illustration of the experimental workflow. NSG-SGM3 mice were injected i.v. with 4 x10^5^ non-senescent or senescent HDF (IR). The following day, 1 x10^5^ A549 tumor cells were injected i.v.. Six days after tumor cell injection, PBMCs and granulocytes (5 x10^6^ cells each/mouse) were injected i.v.. Tumor growth was evaluated every week until day 31, when the mice were sacrificed and their lungs collected for analysis. B) Growth curves for A549 tumor cells injected alone (black) or co-injected with non-senescent HDF (red) or senescent HDF induced by irradiation (blue) in mice lacking hCD45⁺ cells. Each line represents the mean tumor growth ± SEM for 8 mice per group. Statistical analysis was performed using a mixed-effects model, followed by Tukey’s multiple-comparison test. **p < 0.01; ****P < 0.0001. C) Representative bioluminescence images of tumor-bearing mice with or without hCD45^+^ cells. Mice were injected with A549 alone or with non-senescent or senescent HDF. Images were captured on days 6, 16, and 31. D) Tumor growth curves for individual mice injected with hCD45⁺ cells. A549 cells were injected alone (black, n = 5), with non-senescent HDF (red, n = 7), or with senescent HDF induced by irradiation (blue, n = 6). Each growth curve is accompanied by a pie chart illustrating the mean percentage of mice that achieved complete tumor elimination (CE).

**Supplementary Figure 2.**
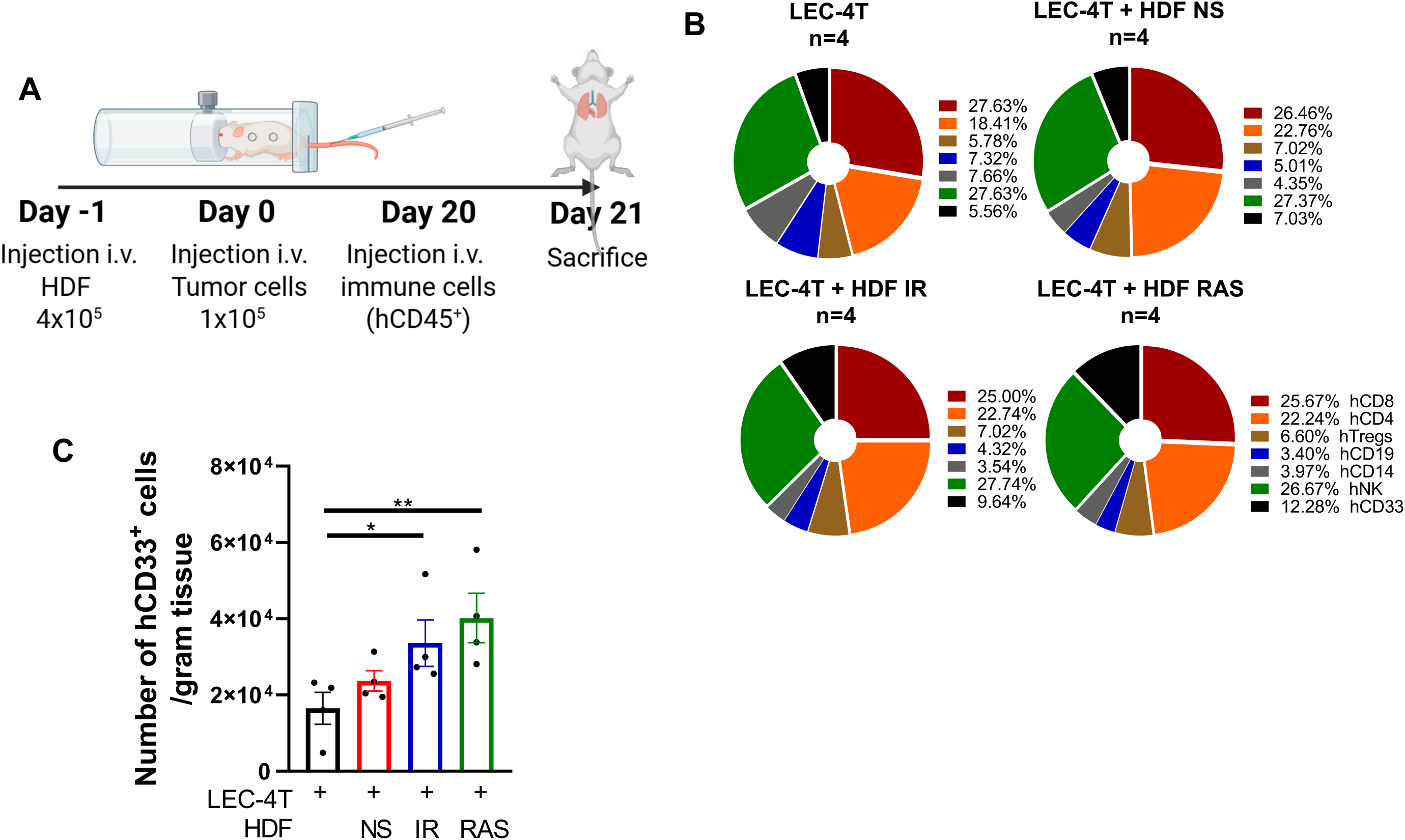
Senescent fibroblasts recruit myeloid cells in tumor-bearing lungs. A. Schematic illustration of the experimental workflow. NSG-SGM3 mice were injected i.v. with 4 x10^5^ non-senescent or senescent HDF induced either by IR or RAS. The following day, 1 x10^5^ LEC-4T tumor cells were injected i.v. and allow to grow for 20 days prior to the i.v. injection of PBMCs and granulocytes (5 x10^6^ cells each/mouse). On day 21, mice were sacrificed and lungs were collected for analysis. B. Pie charts representing the relative frequencies of human immune cell subsets infiltrating the lungs from each indicated group. C. Graph showing the absolute counts of lung-infiltrating hCD33^+^ cells per gram of tissue, determined by flow cytometry. Each dot indicates the count for an individual mouse. Data are presented as mean ± SEM. Statistical significance was assessed using one-way ANOVA followed by Dunnett’s multiple comparisons test. *p < 0.05; **p < 0.01.

**Supplementary Figure 3.**
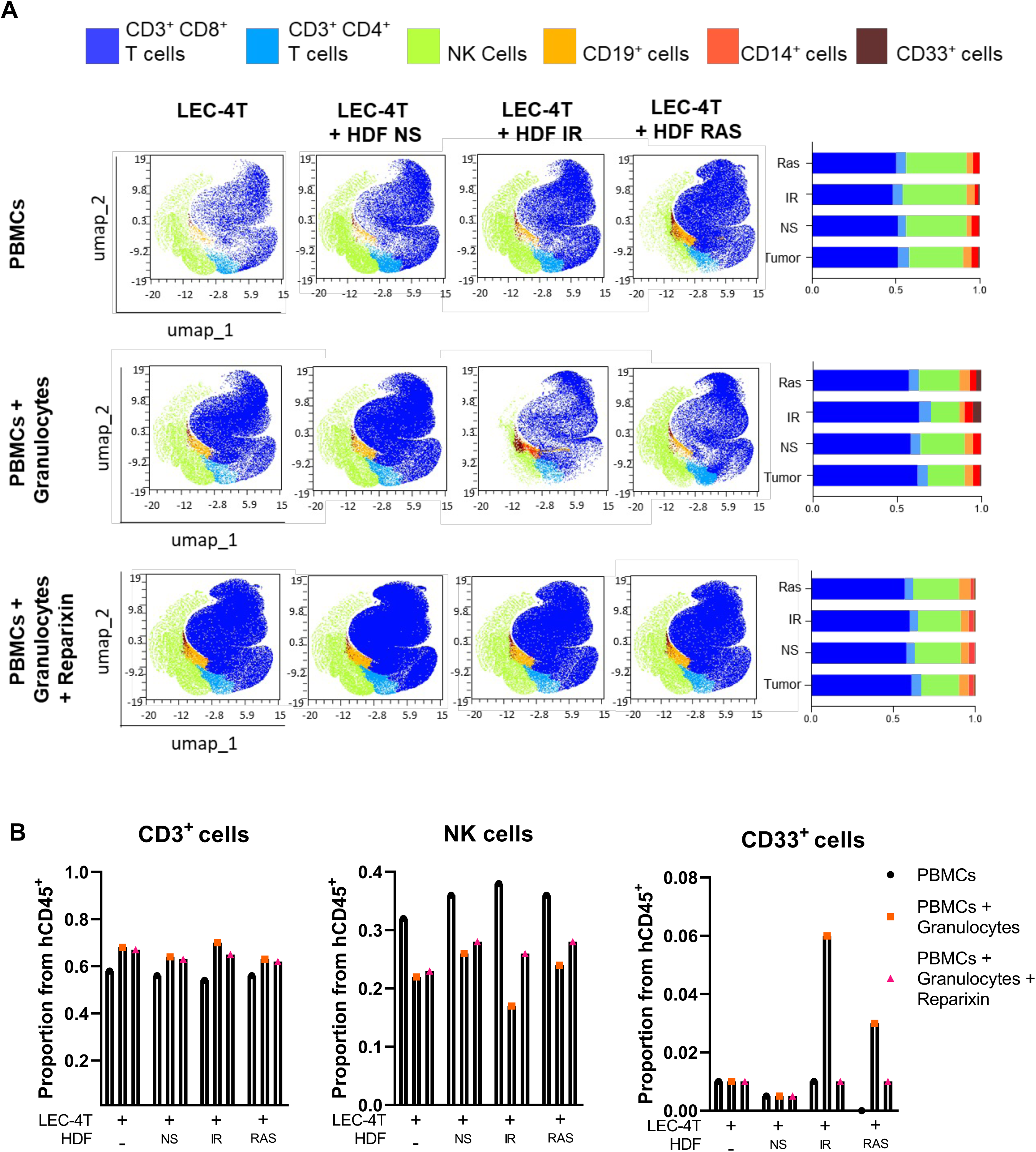
Senescent fibroblasts affect the immune cell subsets infiltrated into tumor spheroids. A. Two-dimensional UMAP plot showing merged clustering of LEC-4T spheroid-infiltrating human immune cell subtypes from Figure 4E. B. Bar graph depicting the frequencies of cell compositions shown in panel A.

**Supplementary Figure 4.**
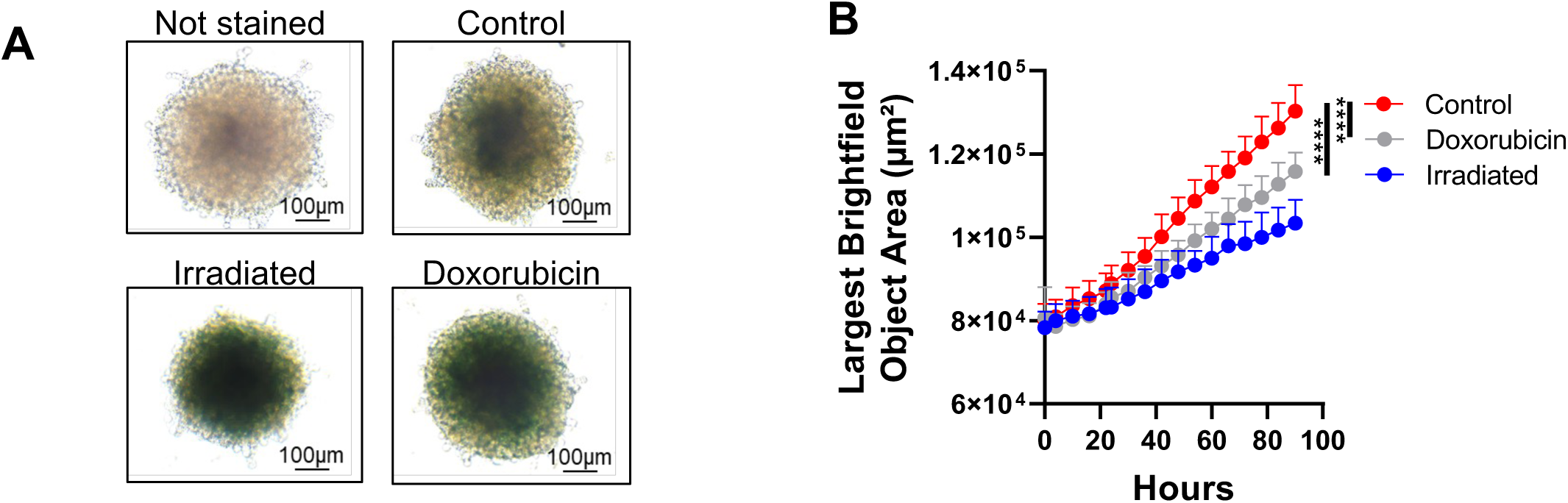
Therapy-induced senescence in spheroids. A. Images of spheroids treated with doxorubicin or irradiation and stained for β-galactosidase activity six days post-treatments. Unstained spheroids and spheroids not exposed to therapy are shown as controls. The scale bar indicates 100 μm. B. Graph representing the average size of spheroids as measured by the largest brightfield object area metric (µm^2^) at various time points post-treatments as measured using IncuCyte imaging. The data represent the mean ± SEM from three biological replicates, each one showing the mean value calculated from 16 spheres per condition. Statistical analysis was performed using mixed-effects modelling with Tukey’s multiple comparisons test. ****P < 0.001.

**Supplementary Figure 5.**
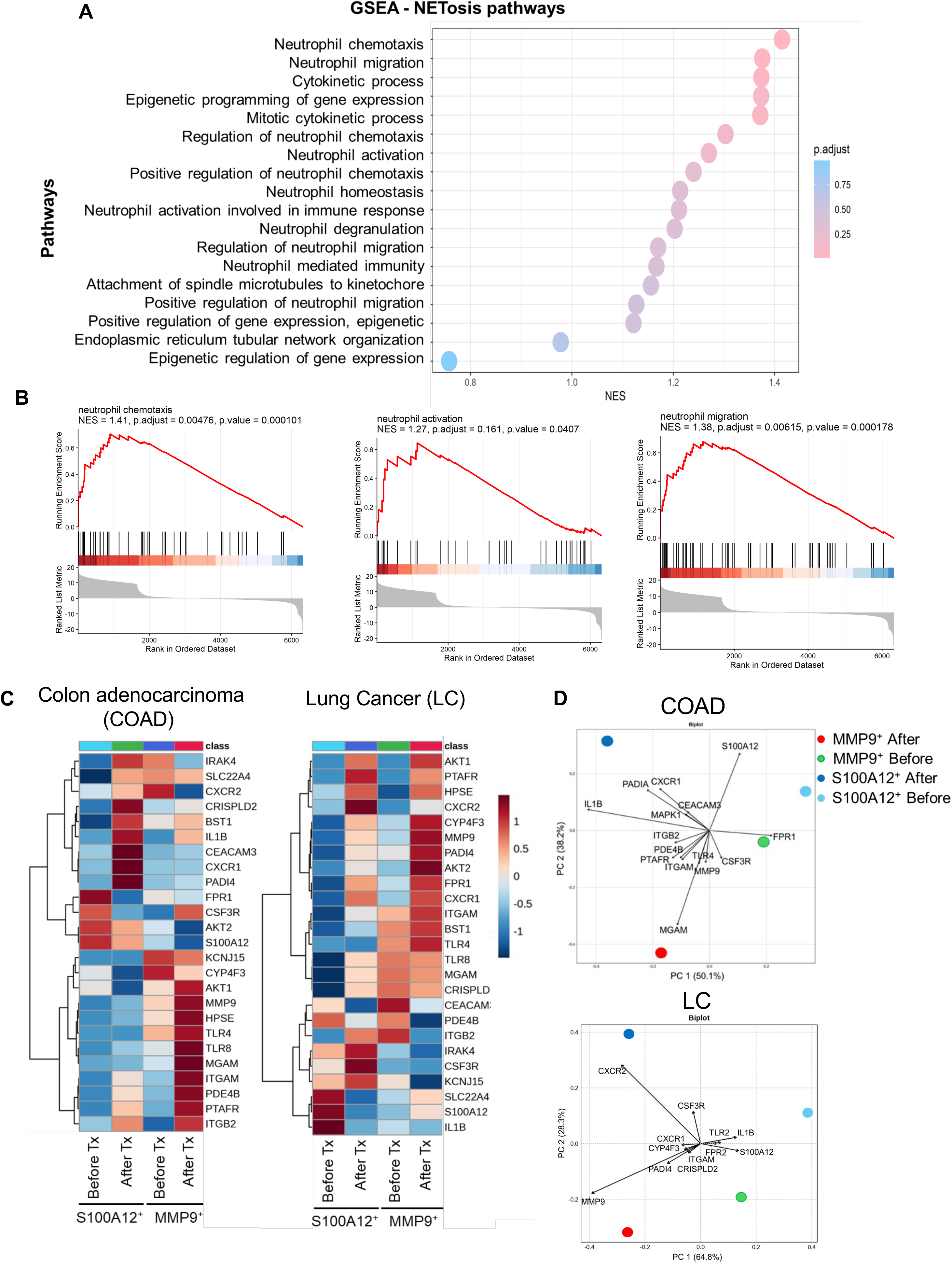
Tumor-infiltrating neutrophils collected from patients before and after genotoxic treatments have increased expression of genes related to NETosis pathways. A. Gene Set Enrichment Analysis (GSEA) of Kyoto Encyclopedia of Genes and Genomes (KEGG) pathways related to NETosis was performed to compare neutrophils from colon adenocarcinoma and lung cancer patient samples before and after genotoxic treatments. The y-axis represents the pathways, while the x-axis indicates their normalized enrichment scores (NES). Adjusted p-values are depicted using color shading. B. GSEA of Gene Ontology categories related to NETosis showed statistically significant enrichment of differentially expressed genes from neutrophil chemotaxis, neutrophil activation and neutrophil activation involved in immune response pathways in patient samples after treatments. C. Heatmaps of colon adenocarcinoma (COAD) and lung cancer (LC) show unsupervised hierarchical clustering of gene sets associated with NETosis in two distinct neutrophil subpopulations (SA10012^+^ and MMP9^+^). The red-to-blue scale represents the log2-transformed fold change value for each gene, with red indicating upregulation and blue indicating downregulation. Each row corresponds to an individual gene, and each column represents samples collected before or after treatments. D. Principal component (PC) analysis biplot of gene expression data, in which neutrophil subpopulations are represented as points and gene expression profiles as arrows. Arrow length reflects the contribution of each variable to the PCs, while the distance between points approximates differences in gene expression patterns among groups. Arrowheads positioned near a specific group indicate that the corresponding genes are expressed at higher relative abundance in those samples.

